# Interactive effect of *Moringa oleifera* mediated green nanoparticles and arbuscular mycorrhizal fungi on growth, root system architecture, and nutrient uptake in maize (*Zea mays* L.)

**DOI:** 10.1101/2025.02.23.639791

**Authors:** Qurat ul Ain, Hafiz Athar Hussain, Lutfur Rahman, Qingwen Zhang, Asma Rehman, Saddam Hussain, Saleem Uddin, Asma Imran

## Abstract

The synergistic interaction between green nanoparticles (NPs) and mycorrhizal fungi promotes plant growth by improving nutrient absorption, optimizing root function, and stress resistance. In this study, we explored the impact of arbuscular mycorrhizal fungus (AMF) *Funnaliformis mosseae* and *Moringa oleifera* mediated green NPs viz; iron oxide (FeO), zinc oxide (ZnO), and Zn doped FeO (Zn/Fe) NPs, both individually and in combination, on maize growth, root system architecture, organic acids production, mycorrhizal colonization, and nutrients uptake. Prior to NP synthesis, metabolomic analysis of *Moringa oleifera* leaves was conducted to characterize its bioactive compounds. The structural properties of green synthesized NPs confirmed by characterization using scanning electron microscopy (SEM), Fourier transform infrared (FTIR) spectroscopy, UV-Vis spectrophotometry, and X-ray diffraction (XRD). The results showed that all individual and combined treatments of AMF and green NPs significantly enhanced maize growth compared to the control. Among the single NP treatments, Zn/Fe NPs exhibited superior performance over FeO and ZnO. However, when combined with AMF, ZnO and Zn/Fe NPs induced the highest growth responses than FeO NPs. The ZnO NPs proved most compatibility with AMF for improving colonization in maize roots, whereas FeO and Zn/Fe NPs reduced colonization efficiency. Furthermore, combination of AMF and ZnO NPs (AMF+ ZnO) noted as most prominent treatment for improvement of maize growth compared to other all treatments, highlighting the potential of AMF+ ZnO NPs for sustainable agricultural applications.

## 1. Introduction

Soil microbes are integral to nutrient cycling and the sustaining soil functions, thereby supporting growth and development of plants (de Moraes et al., 2023; Handa and Kalia, 2024). Moreover, their activities can profoundly modulate plant productivity within agricultural ecosystems, making them critical determinants of crop health and yield (Vincze et al., 2024). Arbuscular mycorrhizal (AM) fungi are among these microbes and are known to have one of the most important symbiotic partnerships in the world (Van Der Heijden et al., 2017). These fungi establish mutualistic interactions with the roots of over 90% of terrestrial plant species (Bonfante and Genre, 2010; van Der Heijden et al., 2015), playing an indispensable role in promoting ecosystem health and plant productivity (Wang et al., 2023).

AM fungi improve plant growth and soil health by aiding in nitrogen fixation, phosphorus solubilization, and producing growth-promoting substances within the rhizosphere (Adedayo and Babalola, 2023). Furthermore, AMF fungi contribute to plant resilience by providing essential nutrients and offering defense against biotic and abiotic, both stresses (Yadav et al., 2023; Hussain et al., 2024). Mycorrhizae are also critical for nutrient exchange in soil, acting as conduits for transferring nutrient from soil to the plants, which, in turn, use photosynthesis to produce carbon. Notably, mycorrhizae can capture up to 15% of the total carbon produced via photosynthesis, functioning as vital carbon sinks in ecosystems (Figueiredo et al., 2021; Hawkins et al., 2023).

The application of plant-mediated green NPs, which make use of organic plant extracts to create NPs in an environmentally friendly way (Khan et al., 2022), is one exciting advancement in agricultural field. However, Moringa oleifera, a highly nutritional and medicinal properties plant (Mahaveerchand and Abdul Salam, 2024), has emerged as promising option for the green NPs synthesis due to its rich phytochemical profile (Barman et al., 2023). These green NPs are distinguished by their biocompatibility, environmentally friendly, and potential to enhance growth and resilience of plants (Khan et al., 2022; Akhter et al., 2024). When combined with AM fungus, these NPs may offer synergistic benefits, improving both the bioavailability of nutrients and plants uptake efficiency, remain relatively less reported. Generally, most crop plant species are responsive to interplay between mycorrhizal symbiosis and NPs (Ingle et al., 2017; Joel et al., 2023).

Maize (*Zea mays L*.), a staple crop with significant importance, faces various challenges related to nutrient availability and environmental stresses exacerbated by the effects of climate change (Shemi et al., 2021; Waqas et al., 2021; Rizwanullah et al., 2023). Addressing these issues through innovative and sustainable practices is imperative for ensuring food security and environmental sustainability (Waqas et al., 2021; Joel et al., 2023). Previous research has showed the efficiency of NPs and AM fungi in alleviating stress and promoting growth in maize (Alsherif et al., 2022; Alsherif et al., 2023; Joel et al., 2023). For instance, the green synthesis of silver NPs (AgNPs) using maize bran-derived arabinoxylans has also been explored as a method to improve maize growth and development, particularly under water-deficit conditions (Raza et al., 2023). Although, previous studies has shown the individual advantages of AM fungi and NPs on plant growth and nutrient uptake, studies exploring their combined synergistic impacts of green NPs with AM fungi in maize are still scarce. So, employing a combined approach of AM fungi and green synthesized NPs to maize plants could prove to be a robust strategy for maize production, especially in regions prone to growth-limiting stresses. To explore this potential, we synthesized three *Moringa oleifera*-mediated green synthesized NPs, viz., ZnO NPs, FeO NPs, and Zn/Fe NPs, and conducted a study to evaluate the effectiveness of green NPs with AMF “*Funneliformis mosseae*” to improve maize growth, root system architecture, mycorrhizal colonization, organic acid profiles in root exudates, and overall nutrient acquisition. We hypothesize that the application of *Moringa oleifera*-mediated green synthesized NPs, either alone or in combination with AMF, has the potential to improve both aboveground and belowground growth, as well as nutrient assimilation in maize plants.

## 2. Materials and methods

### 2.1 Preparation of *Moringa Oleifera* extract

Fresh leaves of *Moringa oleifera* were provided by Department of Agronomy, University of Agriculture, Faisalabad, Pakistan. The leaves were thoroughly washed with tap water and subsequently rinsed with distilled water three to four times to eliminate impurities After washing, the leaves were air-dried in a shaded area at room temperature. Once dried, they were finely powdered using a Silver Crest powder grinder (SC-1880).

For the preparation of the aqueous extract, 10 g of the powdered leaves were dispersed in 90 mL of deionized water and heated at 70°C for 20 minutes. The mixture was then filtered by filter paper Whatman No. 1 to get a purified extract. For subsequent experimental use, the resulting *Moringa oleifera* extract was kept in refrigerator at the 6°C.

### 2.2 Metabolomic analysis of *Moringa Oleifera* leaves

Plant polyphenol metabolomics and targeted metabolome analysis of *Moringa oleifera* leaves powdered were performed using UPLC-ESI-MS/MS system (UPLC, Waters Acquity I-Class PLUS; MS, Applied Biosystems QTRAP 6500+) and UHPLC-MRM-MS/MS by Beijing Biomarker Technologies Co., LTD. The detailed methodology of metabolic analysis is given in Supplementary material.

### 2.3 Synthesis of Moringa oleifera-mediated NPs

Zinc acetate dihydrate (Zn (CH CO) ·2H O; Sigma-Aldrich, 99.99% purity) was used as the zinc ion precursor for the green synthesis of *Moringa oleifera* mediated ZnO NPs. First, 0.2 M zinc acetate stock solution was made in deionized water. To initiate the synthesis, 50 mL of zinc acetate solution was dropwise added to 30 mL of *Moringa oleifera* aqueous leaf extract while being continuously stirred with a magnetic stirrer. A homogeneous solution was obtained by heating the reaction mixture to 70°C. Next, 5.0 M NaOH was added to the mixture dropwise while it was continuously stirred to get its pH to 12. Following the reaction, a yellowish precipitate was formed, indicating the successful ZnO NPs synthesis (Fig. 1). Two or three full washes with deionized water were performed on the precipitate to ensure that any remaining contaminants were eliminated. Finally, the purified product was calcined in a muffle furnace at 200°C for 2 hours to achieve complete crystallization and the elimination of organic impurities. To synthesize *Moringa oleifera* mediated FeO NPs, 0.15 M ferric chloride hexahydrate (FeCl_3_·6H_2_O) solution was added dropwise to *Moringa oleifera* extract in 1:1 volume ratio at 60 °C (Fig. 1). As the FeCl_3_ solution was gradually added to the extract of *Moringa oleifera*, the mixture color turned dark black from a brown color, indicating the synthesis of FeO NPs (Aisida et al., 2020). Furthermore, the reaction mixture was incubated for 4 hours at 60 °C to complete the reaction. In order to eliminate the unreacted precursors, the synthesized NPs were finally obtained by centrifugation at 3500 rpm and repeatedly cleaned with deionized water. After that, the produced NPs were kept at room temperature for later use after being dried for overnight at 60 °C in a vacuum oven.

**Figure 1:**
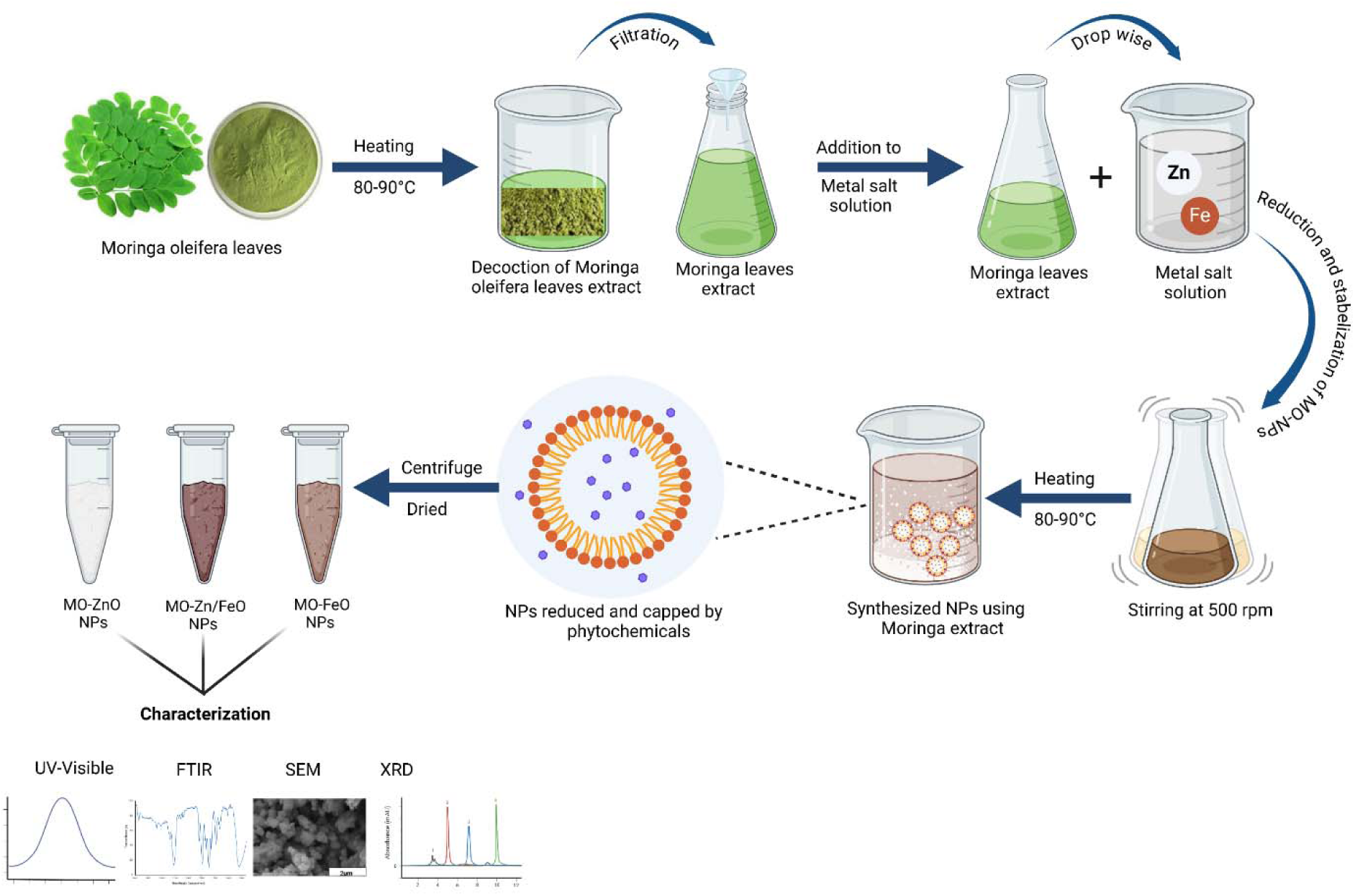
Schematic representation of the green synthesis of *moringa oleifera* mediated green Zinc Oxide (ZnO), Iron Oxide (FeO), and Zn-doped FeO (Zn/Fe) nanoparticles (NPs).

For the synthesis of Zn/Fe NPs, 50 mL solution was prepared containing 1.99g of FeCl_2_.4H_2_O, 5.4g of FeCl_3_.6H_2_O, and 0.8g of Zn (CH_3_CO_2_)_2_.2H_2_O in deionized water. Then, 20ml extract of *Moringa oleifera* leaf was added dropwise to the prepared solution for 30 minutes vigorous stirring at 70°C (Fig. 1). Using 25% NH OH solution, the mixture’s pH was adjusted to 10. The reaction proceeded for two hours at a constant stirring at a temperature of 80 °C (Gholami et al., 2019). After allowing the reaction mixture to naturally cool, the synthesized nanocomposite was cleaned with deionized water until its pH reached neutral. Then, nanocomposites were placed in oven for 2 hours at 150, followed by calcination at 800 for 4 hours to achieve enhanced crystallinity and thermal stability.

### 2.4 Characterization of NPs

The surface morphology of the green NPs was analyzed using a Scanning Electron Microscope (SEM, JSM-6360) at various magnification levels to evaluate particle size and surface texture. The optical properties of the synthesized NPs were analyzed by recording UV-visible spectra using a Shimadzu UV-1700 UV-V is spectrometer (Kyoto, Japan) within a wavelength range of 200–800 nm, employing distilled water as the baseline reference. The structural and crystalline characteristics of the NPs were assessed using a Shimadzu XRD instrument at 50 kV and 20 mA with Cu-Kα radiation (λ = 1.541 Å). In addition, FT-IR spectroscopy (Shimadzu 8400s, range 400–4000 cm ¹) was utilized to pinpoint phytochemical components and functional groups implicated in the stability, capping, and reduction of the synthesized nanomaterials.

### 2.5 Inoculation experiment and growth conditions

A pot experiment was conducted in a controlled greenhouse at the National Institute for Biotechnology and Genetic Engineering (NIBGE) in Faisalabad, Pakistan, during the March to May 2022. The “Zhengdan 958” maize hybrid seeds were planted in earthen pots that were 18 cm in diameter and 20 cm deep, and that were each filled with 3 kg of autoclaved soil. Each pot was initially planted with four seeds, and after seedling emergence, only one healthy plant was kept in each pot to maintain uniform plant density. Soil preparation, fertilization, growth conditions, and irrigation management were conducted following the outlined in our previous study (Ain et al., 2024).

The experiment was designed using a completely randomized design (CRD) consisting of eight different treatments; T1: control (Cn, no NPs or no fungus), T2: arbuscular mycorrhizal fungus (AMF) *Funnaliformis mosseae*, T3: iron oxide (FeO) NPs, T4: zinc oxide (ZnO) NPs, T5: Zn doped FeO (Zn/Fe) NPs, T6: arbuscular mycorrhizal fungus + iron oxide NPs (AMF + FeO), T7: arbuscular mycorrhizal fungus + zinc oxide NPs (AMF + ZnO) and T7: arbuscular mycorrhizal fungus + Zn doped FeO NPs (AMF + Zn/Fe). Each treatment was replicated three times, with three pots in each replicate, for a total of nine plants per treatment.

### 2.6 AMF inoculation and NPs application

Based on the finding of (Hussain et al., 2021), we used *Funneliformis mosseae* (BGC XZ02) for mycorrhizal fungus inoculation in maize at the time of sowing. The source of the mycorrhizal fungus and the method of inoculation were consistent with those discussed earlier (Ain et al., 2024).

Three different NPs were evaluated, either with or without AMF: 100 mg/kg of zinc oxide (ZnO) NPs, 50 mg/kg of iron oxide (FeO) NPs, and 50 mg/kg of zinc-doped iron oxide (Zn/Fe) NPs were tested. Each NP suspension was prepared in deionized water and subjected to sonication for 20 minutes using a Branson Model B200 ultrasonic bath to prevent agglomeration and ensure uniform dispersion.

Seven days after sowing, the nanoparticle suspensions were applied to the soil using a fertigation method, while the control plants received an equivalent volume of deionized water. The maize plants were grown for 45 days, then harvested for morphological and physiological analyses. Additionally, fresh root samples were collected to assess root colonization.

### 2.7 Determination of plant growth characteristics

Plant height was assessed using a calibrated metallic ruler, while stem diameter was determined with a precision digital caliper (Model 500-197-30, Mitutoyo Group, Japan). The total number of leaves per plant was manually counted. Plant height for each pot was recorded individually, and the mean values were calculated to represent the average height per treatment. The plants were collected and divided into aboveground (shoots) and belowground (roots) components after 45 days of growth. While the roots were first completely cleansed with tap water and then deionized water to remove any remaining soil particles, the shoots were carefully washed with distilled water to remove surface pollutants.

Root morphological characteristics, such as total length, volume, average diameter and surface area, were analyzed using an Epson Perfection V700 Photo Flatbed Scanner (B11B178023, Indonesia) and the WinRHIZO. Subsequently, roots and shoots were then oven-dried for 72 hours at 70°C until a consistent weight was reached. The dry weight of both shoots and roots was recorded, and the dried samples were preserved for subsequent nutrient analysis.

### 2.8 Determination of phosphatase (Pase) activity and rhizosheath pH

The activity of phosphatase (Pase) enzyme in the rhizosphere soil was quantified using p-nitrophenyl phosphate (p-NPP) as the substrate (Alvey et al., 2001). Entire root systems with attached soil were carefully shifted into 200 mL of 0.2 mM CaCl solution, with the volume adjusted for each root system according to the protocol outlined by (Veneklaas et al., 2003). To maintain consistency, the pH of the 200 mM Na-acetate buffer was set to 7.4, reflecting the average pH of the rhizosphere samples. After that, rhizosphere soils were separated from the roots through centrifugation at 12,000 × g for 10 minutes, then dried in an oven at 60°C, and subsequently weighed. The spectrophotometer was used to measure the amount of p-NPP in the supernatant at the wavelength of 405 nm.

For rhizosheath pH measurement, maize roots were gently agitated for removing loosely adhering bulk soil. The more firmly attached rhizosheath soil was then carefully collected using a soft brush, ensuring minimal disturbance to the root structure. This soil was suspended in distilled water at 1:5 ratio of soil to water, agitated for 30 minutes, and let to settle down. The pH of the solution was precisely determined with a calibrated pH meter.

### 2.9 Estimation of AMF colonization in roots

To calculate AMF root colonization, root segments (1 cm in length, 30 segments used as replicates per sample) were processed using a sequential staining approach in order to calculate AMF root colonization. After 1.5 hours of clearing the roots in a 10% (w/v) KOH solution at 95°C, the roots were bleached for 15 minutes in a 10% (v/v) H_2_O solution. Following bleaching, the roots were acidified in 0.2 M HCl for an hour, and they were then stained for three minutes with 0.05% (w/v) trypan blue in a lactoglycerol solution (Phillips and Hayman, 1970). Using a light microscope at 200–400x magnification, the magnified gridline intersect methods was used to quantify root colonization in accordance with the protocol described by (Giovannetti and Mosse, 1980). Mycorrhizal dependency (MD) measure of how plants respond to AMF inoculation in terms of growth as opposed to non-inoculated plants, was calculated using the following equation (1) (Tawaraya and Nutrition, 2003; Van Der Heijden et al., 2003):

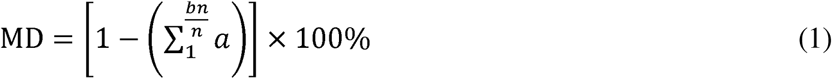

where a represents the dry mass of plants inoculated with AMF, b is the dry mass of non-inoculated plants, and n is the number of replicates. This metric provides a quantitative measure of the extent to which mycorrhizal inoculation enhances plant growth relative to non-mycorrhizal treatments.

### 2.10 Determination of organic acids from rhizosheath soil

The organic acid concentration in the rhizosheath soil was determined by HPLC in ions suppressing mode, according to instructions by (Szmigielska et al., 1996). Initially, 10 g of rhizospheric soil was extracted with 50 mL of ultrapure water and subjected to vigorous shaking for two hours to maximize the solubility of organic acids. Following a 15-minute centrifugation at 8,000 x g, the suspension was filtered through a 0.45 μm membrane to eliminate any remaining particles from the supernatant. To isolate the organic acid anions, the filtered supernatant was passed sequentially through cation and anion exchange resin columns, and the bound anions were subsequently eluted with 1 M HCl. The eluate was concentrated using a rotary evaporator set at 90 rpm until the volume was reduced, after which it was diluted back to 10 mL with ultrapure water. A final filtration step using a 0.45 μm filter ensured the purity of the extract before HPLC analysis. Organic acids were noted by UV absorption at 214 nm, utilizing a mobile phase of 25 mM KH PO buffer (pH 2.25), the HPLC analysis was carried out on a 250 mm × 4.6 mm reversed-phase C18 column at a temperature of 31°C and a flow rate of 1 mL/min. The organic acids were identified and quantified through comparison of the retention durations of the sample peaks to those of established standards, including citric, fumaric, malonic, malic, oxalic, and trans-aconitic acids. Calibration curves were generated for each standard, and the concentrations of the organic acids were represented on the basis of dry soil to ensure accuracy and comparability across samples.

### 2.11 Determination of nutrient concentration and uptake

Following oven drying the plant samples, 300 mg of samples were finely crushed into powder and digested in a sealed chamber using 8 mL of concentrated sulfuric acid (H SO) (Anderson and Henderson, 1986). Using the Kjeldahl method, nitrogen was measured as outlined by (Sivasankar and Oaks, 1995). Elemental Analyzer was used to measure the potassium, while UV/visible spectrophotometer was used to quantify phosphorus using the vanadate-molybdate method according to the protocol by (Chapman and Pratt, 1962). Nutrient uptake was estimated by using the Equ. (2) as follows,

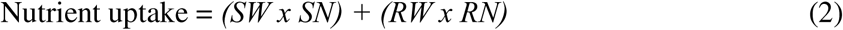

where SW represents the shoot dry weight (g), SN is the nutrient concentration in shoots (g kg ¹), RW denotes the root dry weight (g), and RN is the nutrient concentration in roots (g kg ¹).

### 2.12 Statistical analysis

The data were statistically evaluated using one-way analysis of variance (ANOVA) in Statistix

8.1 (Analytical Software, Tallahassee, FL, USA). Differences among treatment means were evaluated using Tukey’s test at a significance level of p < 0.05. Graphical visualizations of the results were generated using SigmaPlot 14.0 and OriginPro 2019b. To investigate complex interactions between variables, SmartPLS 4 was used for partial least squares path modeling

(PLS-PM). Correlation analyses were performed using the corr.test() function in the R statistical software to investigate relationships between various parameters.

## 3. Results

### 3.1 Polyphenols and targeted metabolomics of *Moringa oleifera*

Total of 354 polyphenols were detected in the qualitative and quantitative analysis of polyphenol metabolome in moringa sample. The composition of metabolites was as, flavonoids 55.65% and polyphenols 44.35%. (Fig S1). The metabolite classification of *Moringa oleifera* was analyzed using three major databases: KEGG, HMDB, and LIPID MAPS, revealing a diverse metabolic composition (Fig 2). In the KEGG database, the predominant metabolite categories were lipids and lipid-like molecules, organic acids, and amino acid-related compounds. The HMDB classification provided a more detailed breakdown, highlighting the abundance of lipids, organic acids, and amino acids, along with a noticeable presence of bioactive secondary metabolites such as phenolics and alkaloids. The LIPID MAPS classification further refined the lipid composition, identifying glycerophospholipids as the most dominant class, followed by sphingolipids, fatty acids, and sterol lipids. Collectively, these results demonstrate that *Moringa oleifera* is metabolically rich in primary metabolites essential for physiological functions, while also containing secondary metabolites that may contribute to its medicinal properties. The detailed information on all identified metabolites and targeted metabolites information is provided in supplementary data file.xlsx.

**Figure 2:**
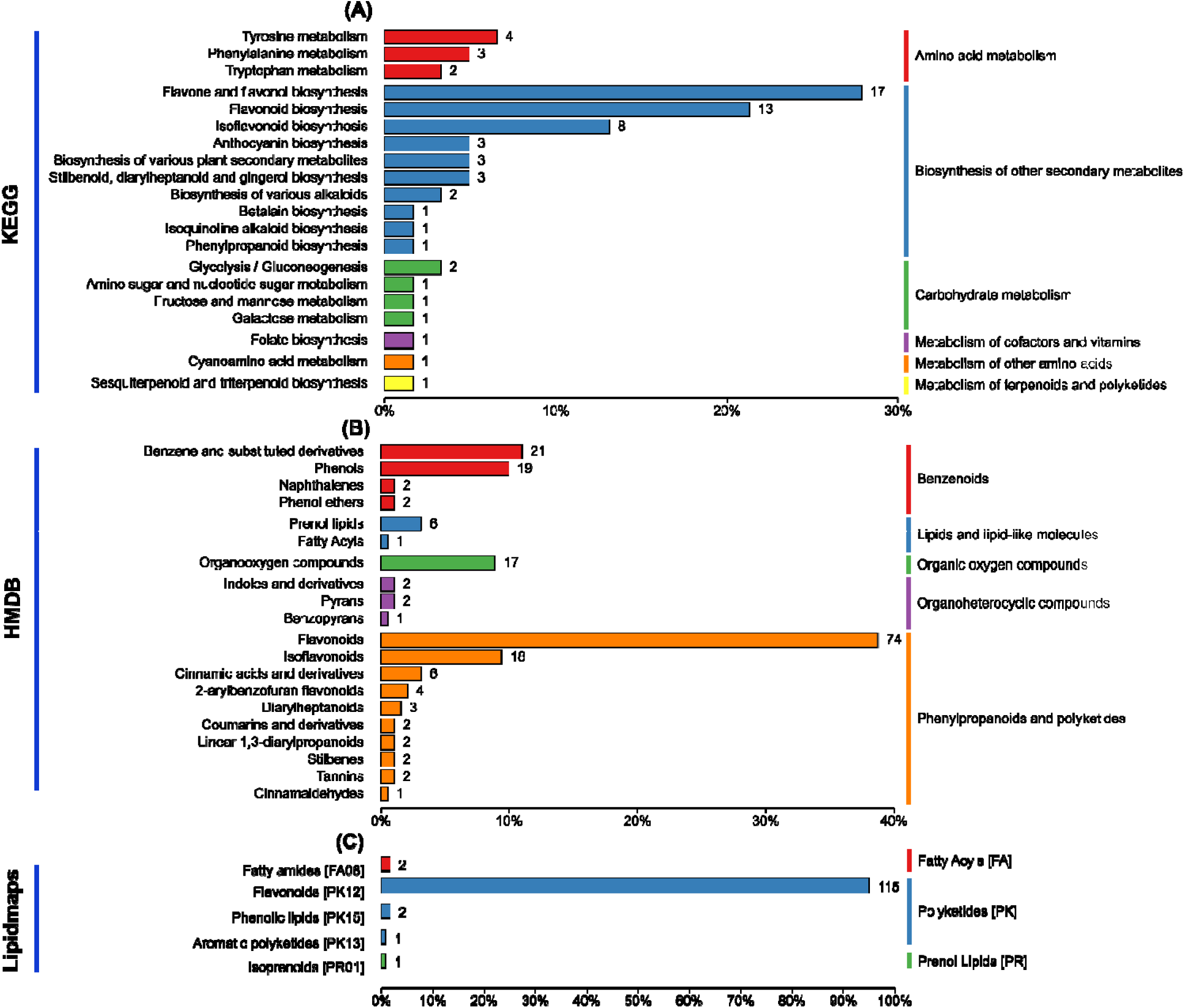
Classification of metabolites based on different databases. (A) Summary of metabolite classification according to the KEGG database. (B) Summary of metabolite classification based on the HMDB database. (C) Classification of metabolites using the LIPID MAPS database. The length of each bar represents the number of metabolites assigned to each category in the respective database. Different colors indicate distinct metabolite classes.

### 3.2 Characterization of NPs

The SEM images show that ZnO NPs have an irregular nanoplate morphology with a high surface area, ideal for catalytic applications, and a uniform size distribution centered around

21.13 nm. In contrast, FeO NPs exhibit a denser, more compact structure with a wider size distribution, averaging 26.80 nm, likely due to different synthesis conditions or material properties. The Zn-doped FeO NPs present larger, more agglomerated structures with an average size of 33 nm and a broader distribution, indicating that Zinc doping impacts both particle size and agglomeration (Fig. 3). The irregular shapes and varying sizes of these composite nanoparticles imply a spontaneous synthesis mechanism, potentially leading to diverse applications based on their unique structural characteristics.

**Figure 3:**
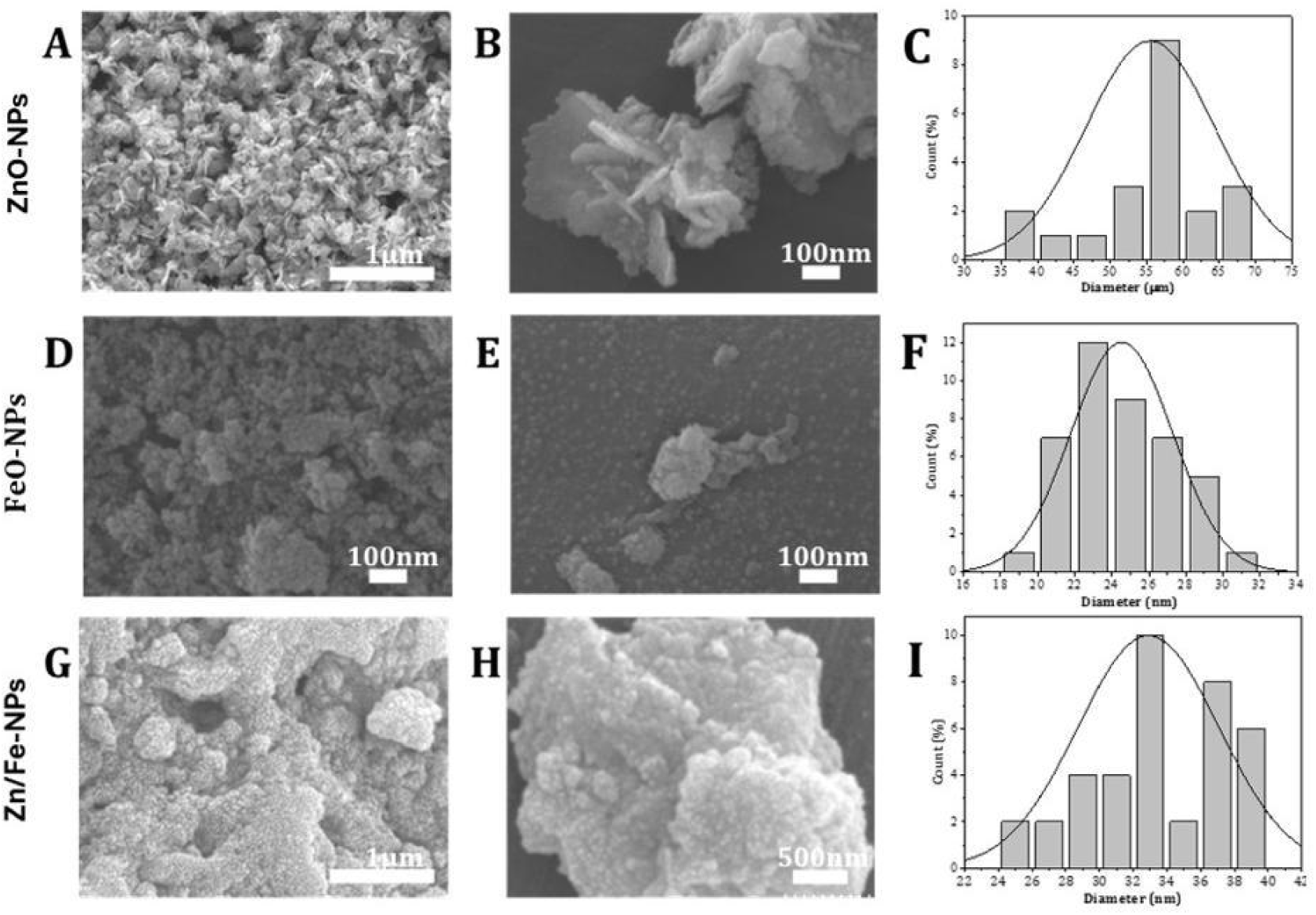
Scanning Electron Microscopy (SEM) analysis of green synthesized nanoparticles (NPs): (A-B) ZnO NPs, (C) size distribution of ZnO NPs, (D-E) FeO NPs, (F) size distribution of FeO NPs, (G-H) Zn/Fe NPs, and (I) size distribution of Zn/Fe NPs.

The zeta potential values of ZnO-NPs, FeO-NPs and Zn dopped FeO-NPs were −33.6 mV ± 6.01 mV, −34 mV ± 7.68 mV and −18 mV ± 5.96 mV respectively (Fig. 4). The analysis showed a singular, sharp peak for each type of nanoparticle with peak area ratios recorded as 17.9 mV/100.0%, 33.6 mV/100.0%, and 33.8 mV/100.0% respectively. The observed negative zeta potentials suggest that the nanoparticles are likely stabilized by bioorganic molecules from plant extracts, which confer a negative charge to their surfaces (Edison and Sethuraman, 2012). These high negative values indicate strong electrostatic repulsions between particles, promoting stability and preventing agglomeration (Sivaraman et al., 2013). Notably, Zn-doped FeO-NPs demonstrated the highest zeta potential (−34 mV), suggesting superior colloidal stability due to enhanced electrostatic repulsion among the particles.

**Figure 4:**
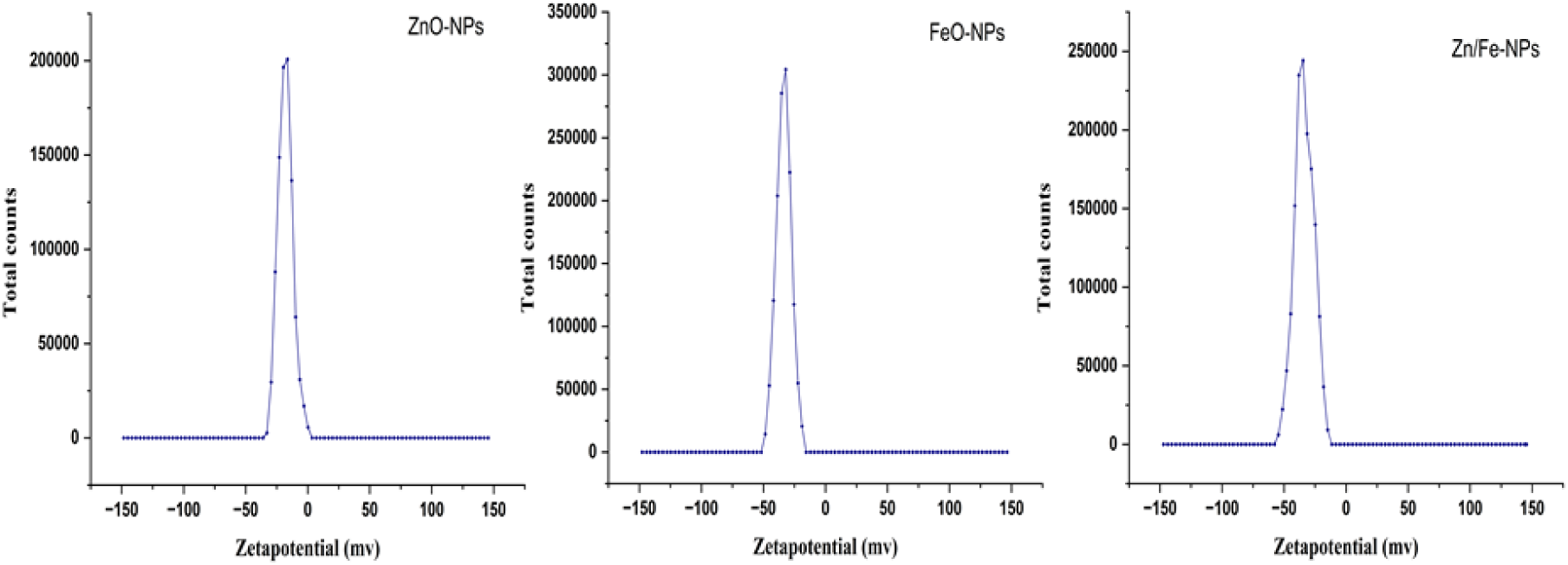
Zeta potential analysis of green-synthesized ZnO, FeO, and Zn/Fe NPs.

The synthesis of NPs was evidenced by characteristic UV-Vis absorption bands observed between wavelengths of 300–800 nm (Fig. 5). The surface plasmon resonance (SPR) bands, which are indicative of the particles’ electronic structure, vary according to the nanoparticles’ shape, size, morphology, dielectric environment, and chemical composition (Liaqat et al., 2022). The SPR bands were recorded with peak absorbance at 341 nm for ZnO-NPs, at 299 nm for FeO-NPs, and at 348 nm for Zn-doped FeO-NPs. It is important to note that the absorption peak for FeO-NPs typically ranges between 280 and 420 nm, aligning with the observed peak at 299 nm, which falls well within this expected range (Pindiga et al., 2022). The slight shifts in peak absorbance observed for ZnO-NPs and Zn-doped FeO-NPs could be attributed to changes in nanoparticle composition and structural modifications induced by zinc doping.

**Figure 5:**
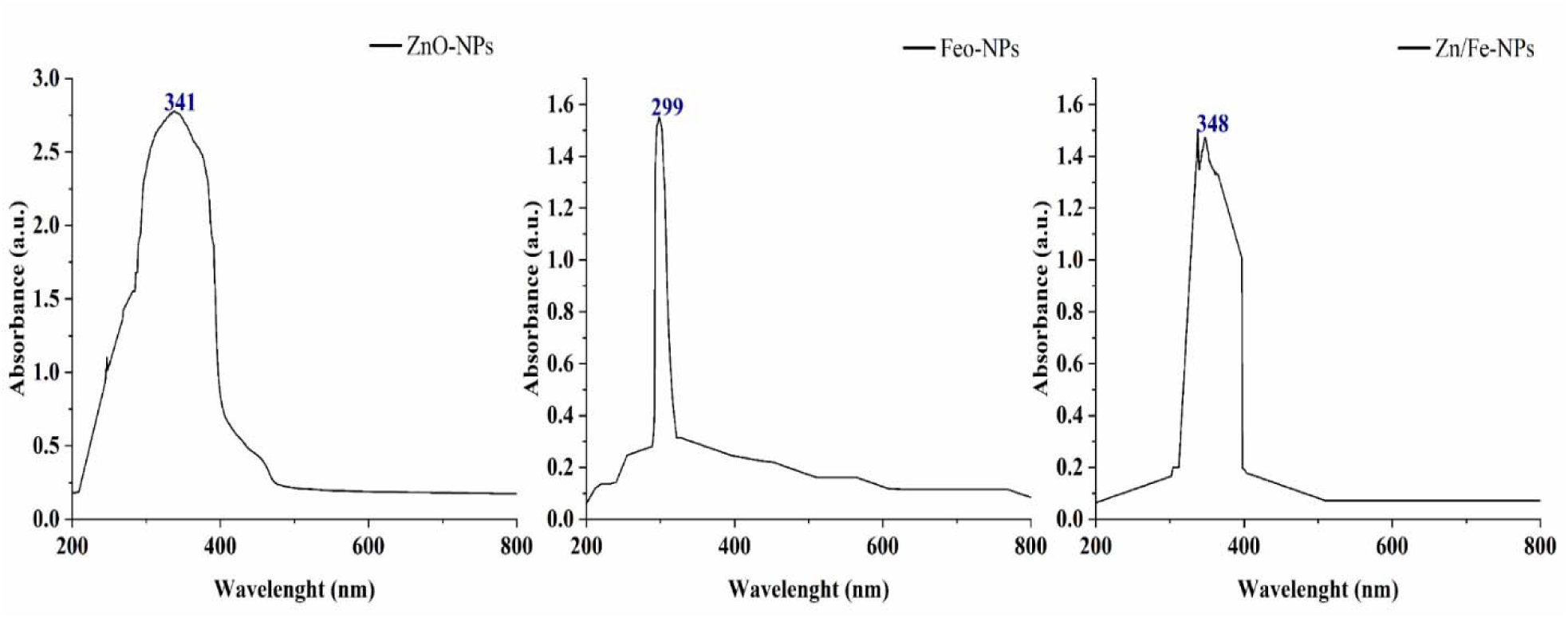
UV-visible spectroscopy analysis of ZnO, FeO, and Zn/Fe NPs. Spectral peaks indicate the characteristic absorbance properties of each NPs type.

X-ray diffraction (XRD) analysis was performed to evaluate the phase purity, crystalline structure, and overall crystallinity of the synthesized nanoparticles. The XRD patterns of ZnO nanoparticles (ZnO NPs) exhibited seven distinct diffraction peaks at 2θ values of 31.68°, 34.34°, 36.16°, 47.48°, 56.54°, 63.10°, and 67.84°, corresponding to the (100), (002), (101), (102), (110),

(103), and (112) crystallographic planes, respectively (Fig. 6). These diffraction peaks are characteristic of the hexagonal wurtzite structure of ZnO, as confirmed by the Joint Committee on Powder Diffraction Standards (JCPDS, card No. 89-7102), indicating the high crystallinity of the synthesized NPs (El-Belely et al., 2021). The average crystallite size was calculated using the Debye-Scherrer formula, centered on the most intense (101) peak at a 2θ value of 36.16°, and was estimated to be approximately 41 nm.

**Figure 6:**
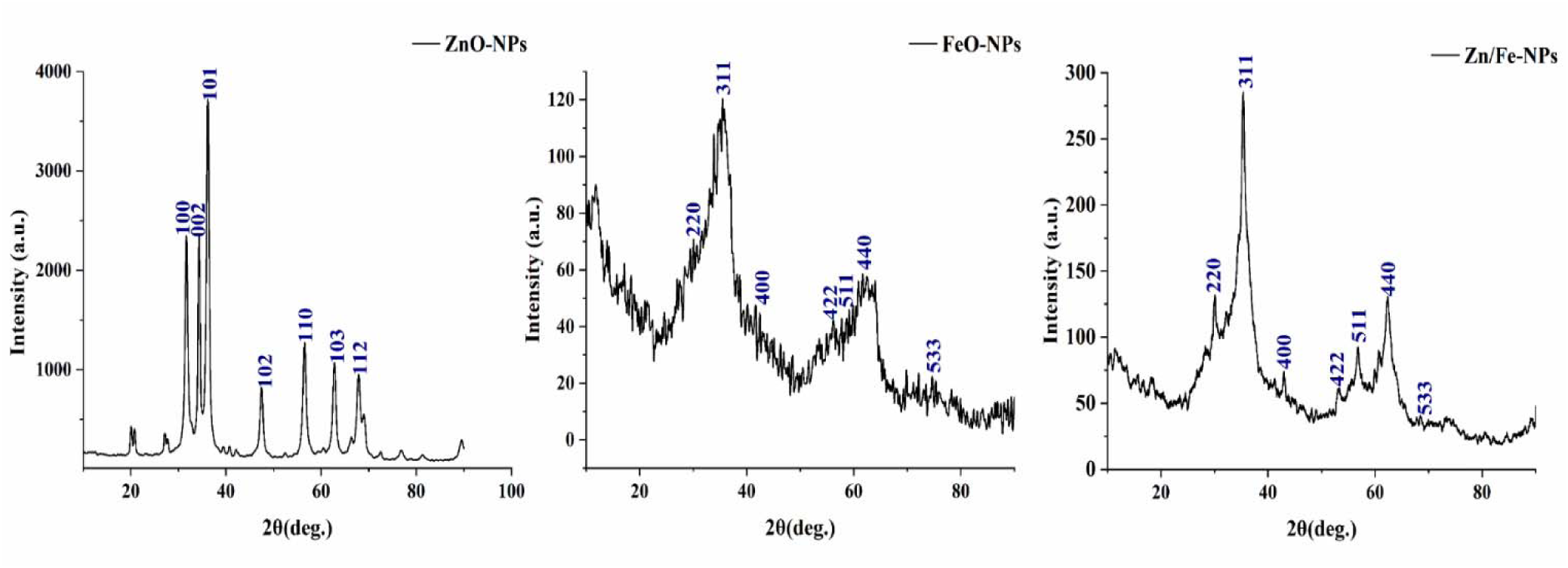
X-ray diffraction (XRD) patterns of ZnO, FeO, and Zn/Fe NPs. The x-axis represents the diffraction angle (2θ) in degrees, while the y-axis represents the intensity in arbitrary units (a.u.).

For the FeO NPs, the XRD pattern displayed prominent peaks at 2θ values of 30.56°, 34.86°, 42.46°, 56.01°, 57.72°, 62.90°, and 74.90°, corresponding to the (220), (311), (400), (422), (511), (440), and (533) planes, respectively. These peaks align well with the standard magnetite (Fe O) structure recorded in the JCPDS file No. 00-003-0863, confirming a cubic crystallographic system (Yew et al., 2016). The sharpness and intensity of the peaks further indicate a well-defined crystalline phase.

For the Zn/Fe NPs, the XRD pattern predominantly exhibited the characteristic peaks of the magnetite phase, with diffraction peaks appearing at 2θ values of 30.06°, 35.68°, 43.40°, 53.84°, 57.44°, 62.96°, and 74.22°, corresponding to the (220), (311), (400), (422), (511), (440), and (533) crystallographic planes, respectively. The retention of these peaks, along with minor shifts, is indicative of successful Zn doping into the Fe O lattice without disrupting its cubic structure, consistent with recent findings (Kasparis et al., 2023). These results validate the successful incorporation of Zn into the magnetite matrix, preserving the overall crystalline framework while potentially enhancing its physicochemical properties.

The Fourier transforms infrared (FT-IR) spectra were used to identify the functional groups in Moringa oleifera extract and produced NPs, as shown in (Fig. 7). The spectrum of the *Moringa oleifera* extract exhibited prominent absorption bands at 3312, 2146, 1641, 1199, and 1095 cm ¹, indicative of several phytochemicals such as amino acids, alkaloids, flavonoids, and phenolics (Moyo, et al. 2012). Specifically, the bands at 1041 and 1199 cm ¹ correspond to C–O stretching vibrations typically associated with alcohols, while the sharp peak at 1641 cm ¹ is attributed to C=O stretching of polar compounds. Additionally, peaks at 1375 and 1641 cm ¹ indicate C=C stretching within aromatic rings, confirming the presence of aromatic structures. For ZnO NPs, the spectral region between 400 and 600 cm ¹ is characteristic of metal–oxygen bonds, with a distinct Zn–O stretching vibration observed at 616 cm ¹ (Karam and Abdulrahman, 2022). Peaks at 2910 and 3238 cm ¹ correspond to O–H stretching vibrations, likely arising from water molecules adsorbed on the nanoparticle surfaces (Kaningini et al., 2022). The band at 1449 cm ¹ is attributed to C–H bonding in alkene structures (Adam et al., 2021), suggesting the presence of hydrocarbon residues.

**Figure 7:**
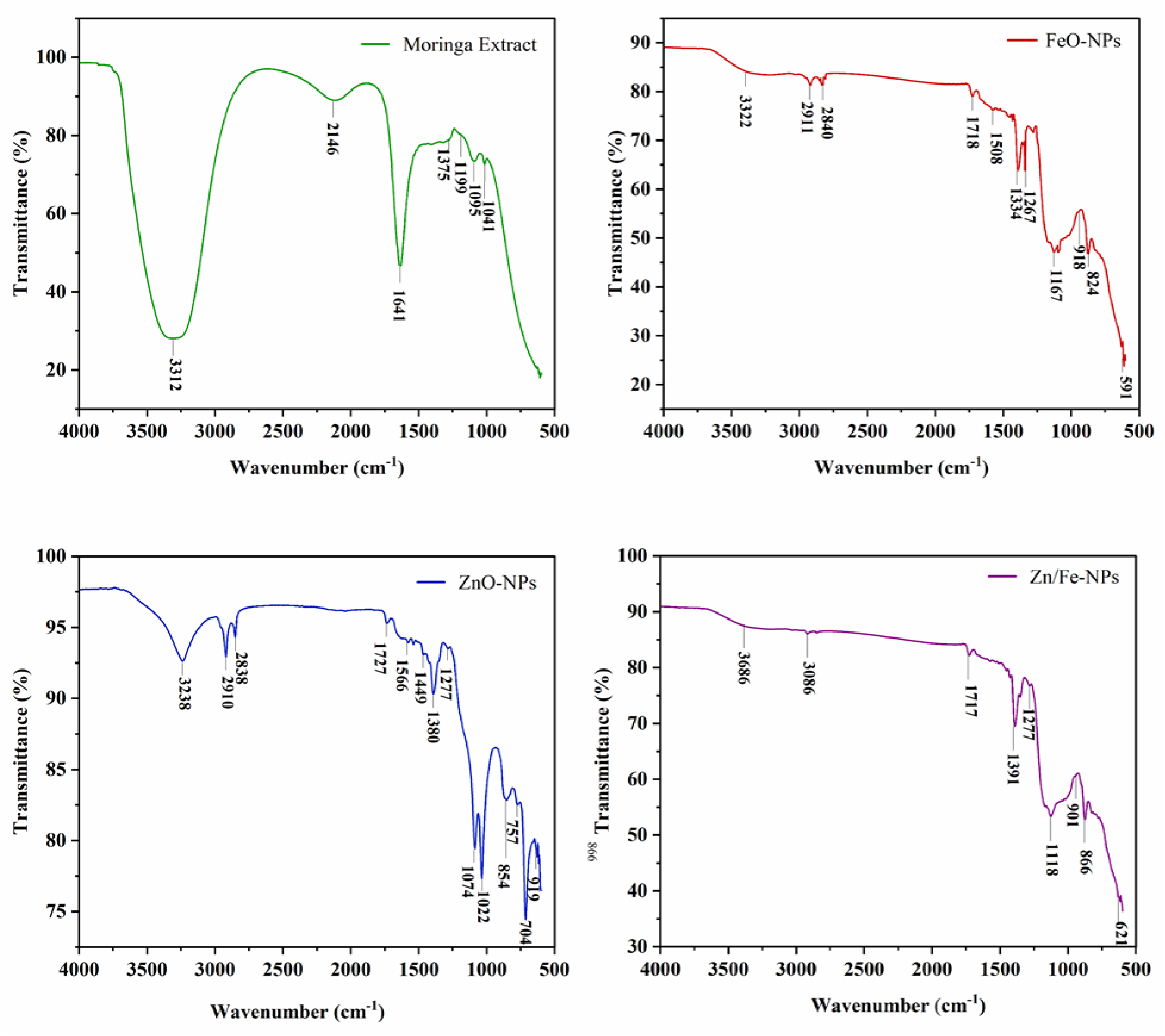
Fourier transforms infrared (FT-IR) spectra analysis of ZnO, FeO, and Zn/Fe NPs.

The FT-IR spectra of FeO NPs and Zn/Fe NPs revealed absorption bands at 3322, 2911, 2840, 1718, 1508, 1334, 1267, 1167, 918, 824, and 591 cm ¹ for FeO NPs, and at 3686, 3086, 1717, 1391, 1277, 1118, 901, 866, and 621 cm ¹ for Zn-doped FeO NPs. The strong absorption bands at 1718 and 1717 cm ¹ are indicative of C=O stretching vibrations, which are associated with polyphenolic compounds and amino acids that act as capping and stabilizing agents during nanoparticle synthesis (Bhuiyan et al., 2020). The bands at 1391 and 1334 cm ¹ correspond to the bending vibrations of O–H groups in phenolic structures (Qasim et al., 2020), further confirming the involvement of phenolic compounds in nanoparticle stabilization.

The FeO NPs spectrum displayed bands at 616 and 704 cm ¹, corresponding to the Fe–O stretching vibration modes within the crystalline lattice of FeO NPs (Aisida et al., 2020). Similarly, the Zn doped FeO NPs spectrum showed absorption bands at 621 cm ¹, attributed to the Zn–O stretching vibrations in tetrahedral sites and Fe–O bonds in octahedral sites of the doped nanoparticles (Lakshmi Ranganatha et al., 2020; Sarala et al., 2020)

### 3.3 Interactive effect of AMF and NPs on growth and biomass allocation of maize

Single and interactive application of AMF and green synthesized ZnO, FeO, and Zn/Fe NPs improve the maize above- and below-ground growth characteristics as shown in (Fig. 8). Among single treatments, Zn/Fe NPs improve fresh and dry weight of shoot, higher plant height and shoot diameter, than AMF, Zn NPs, and FeO NPs, as compared with control treatment (no AMF or NPs). However, a higher no. of leaves and leaf area was noted in AMF treatment than in single treatments of NPs. In interactive treatments, AMF + ZnO-NPs improves higher shoot growth than AMF+ FeO NPs and AMF+ Zn/Fe NPs, as compared with control. Overall, shoot growth were most significantly (P ≤ 0.05) increased under AMF + ZnO NPs treatment.

**Figure 8:**
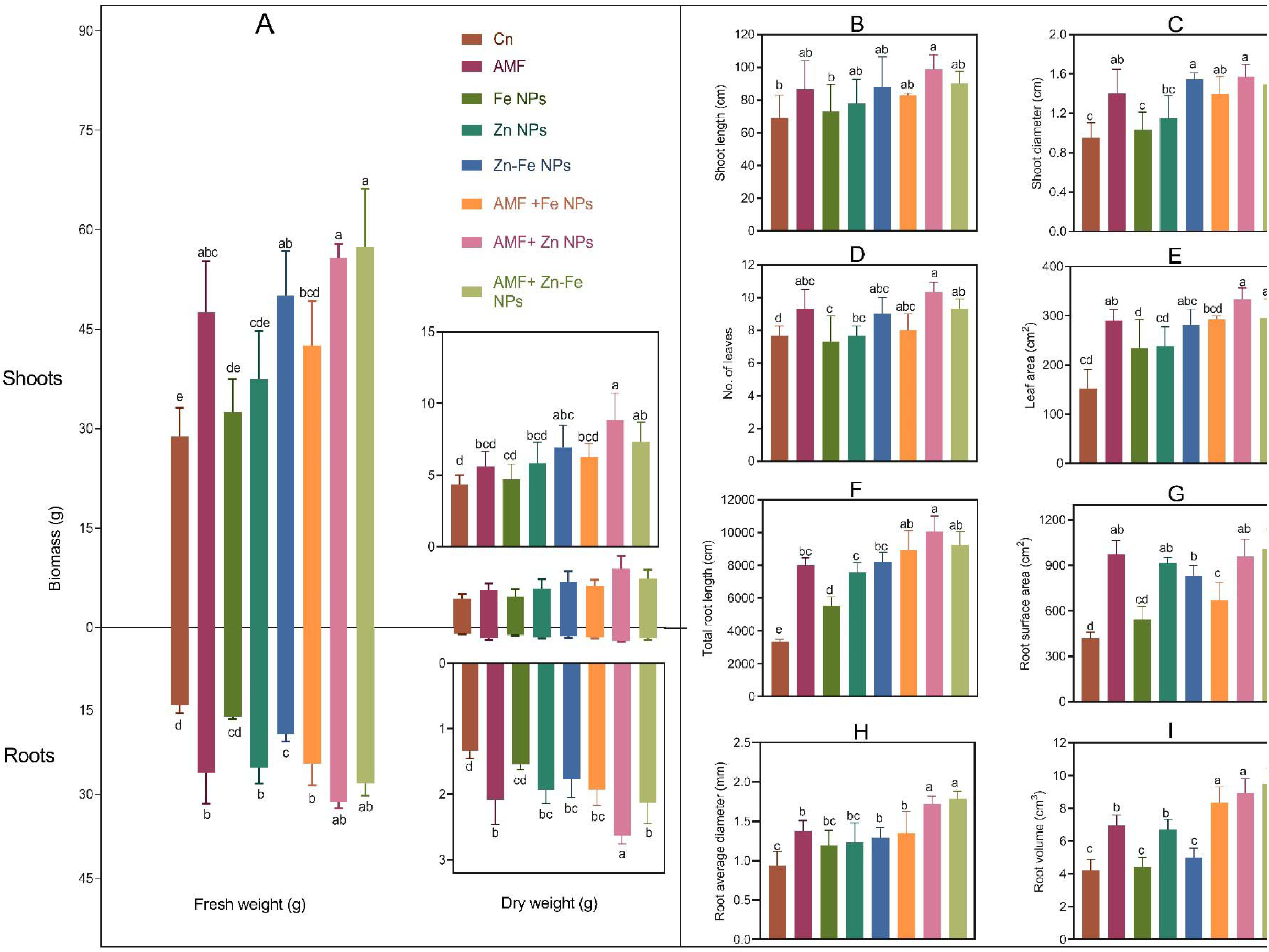
Interactive effect of NPs and AMF on (A) biomass (g) (shoot fresh weight, shoot dry weight, root fresh weight, root dry weigh), (B) shoot length, (C) shoot diameter, (D) no. of leaves, (E) leaf area, (F) total root length, (G) root surface area, (H) root average diameter and (I) root volume of the maize. Error bars represent ± SE of three replicates. Different letters above the bars indicate statistically significant differences among treatments (p < 0.05).

Application of AMF and green synthesized NPs significantly (*P* ≤ *0.05*) improve the growth of maize roots as shown in (Fig. 8). Among the individual treatments, AMF exhibited superior performance in increasing fresh and dry weight of roots, total root length, average root diameter, root surface area, and root volume compared to FeO, ZnO, and Zn/Fe NPs, relative to the control. In combined treatments, the AMF + ZnO NPs combination was notably effective in improving total root length, root fresh weight, and root dry weight, compared to AMF + FeO NPs and AMF + Zn/Fe NPs, relative to the control. Conversely, the greatest increases in average root diameter, root volume, and root surface area were observed with the AMF + Zn/Fe NPs treatment, surpassing the enhancements seen with AMF + FeO NPs and AMF + ZnO NPs, when compared to the control.

### 3.4 Interactive effect of AMF and NPs on phosphatase activity and rhizosheath pH

Data for the individual and interactive effect of AMF and different green synthesized NPs on phosphatase activity and rhizosheath pH was presented in (Fig. 9). Among single treatments, Zn/Fe NPs improve higher pase activity than AMF, FeO NPs and ZnO NPs treatments, as compared with control. Interestingly, FeO NPs decreases the pase activity, as compared with control. Under interactive treatments, AMF + ZnO NPs increase higher pase activity than AMF + FeO NPs and AMF + Zn/Fe NPs, as compared with control. However, non-significant (*P* ≥ *0.05*) difference was noted among the single and combine treatments of AMF and NPs for rhizosheath PH, as compared with the control.

**Figure 9:**
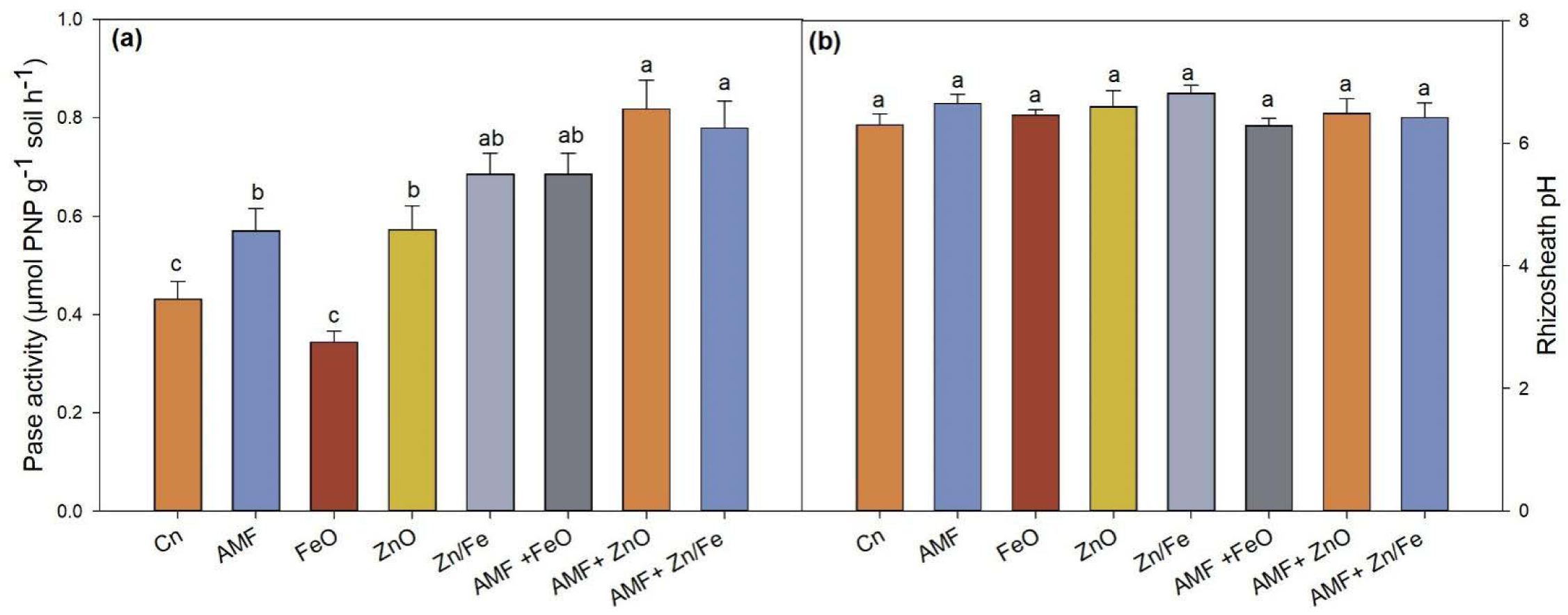
Interactive effect of NPs and AMF on Pase activity and soil pH. Error bars represent ± SE of three replicates. Different letters above the bars indicate statistically significant differences among treatments (p < 0.05).

### 3.5 Effect of NP on AMF colonization and mycorrhizal dependency

The data for effect of green synthesized NPs on mycorrhizal colonization and dependency is presented in (Fig. 10). For mycorrhizal colonization, AMF+ZnO NPs was the best treatment to improve colonization but AMF+FeO and AMF+Zn/Fe decreased mycorrhizal colonization as compared with the treatment without NPs (AMF). However, there was no statistically (*P* ≥ *0.05*) difference among treatments for root colonization. In contrast, mycorrhizal dependency was decreased by FeO NPs, compared with the treatment without NPs. The mycorrhizal dependency was most increased in ZnO NPs, compared with AMF, AMF +FeO NPs and AMF + Zn/Fe NPs. The significant (*P* ≤ *0.05*) improvement of mycorrhizal dependency was noted in ZnO NPs and Zn/Fe NPs.

**Figure 10:**
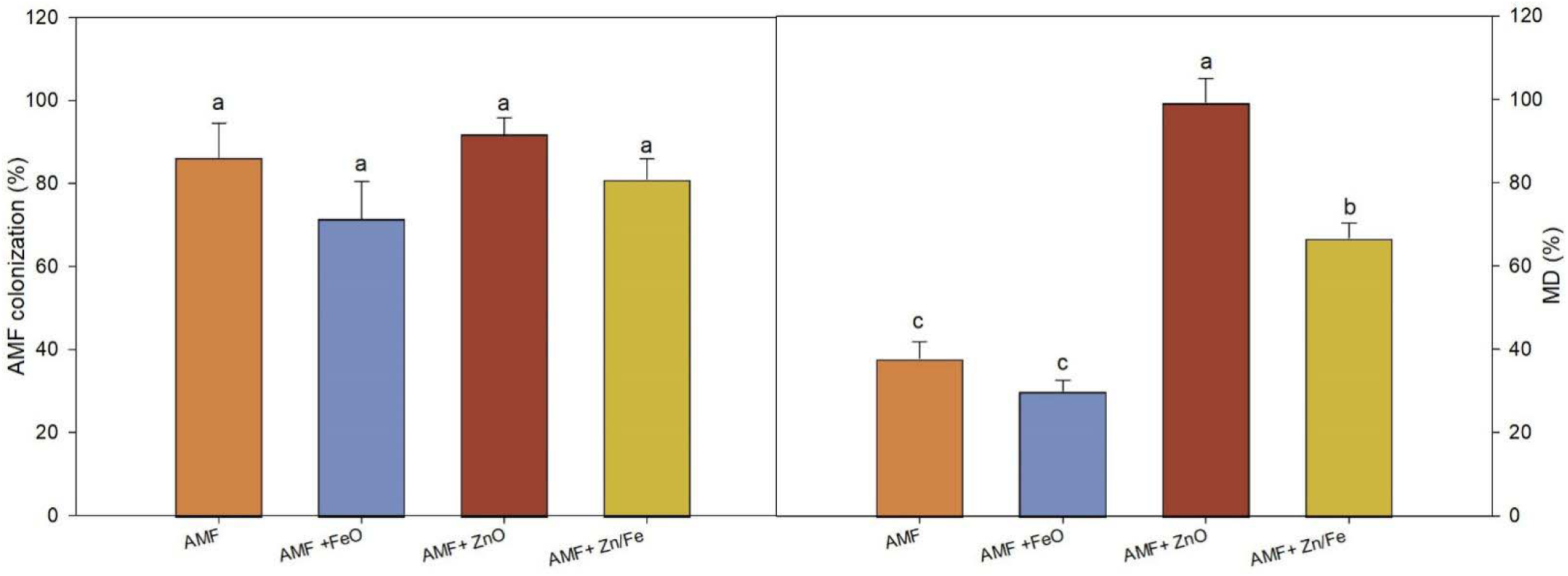
Effect of NPs on mycorrhizal colonization and mycorrhizal dependency (MD) percentage (%). Error bars represent ± SE of three replicates. Different letters above the bars indicate statistically significant differences among treatments (p < 0.05).

### 3.6 Interactive effects of NPs and AMF on the organic acids in maize roots

The data for individual and interactive effects of AMF and green synthesized NPs on organic acid of maize roots is given in Table 1. Individual AMF treatment increased the production of oxalic, citric and malic, while lactic, T-aconitic and succinic acids were reduced by AMF as compared to the control treatment. The treatments with individual NPs generally increased organic acid production than the AMF treatment. Among all individual treatments, higher production of oxalic and malic were recorded under AMF treatment, T-aconitic was noted under ZnO NPs treatment, citric, lactic and succinic were noted under Zn/Fe NPs treatment, as compared with control. The interactive treatments of AMF and NPs often led to higher organic acid production than AMF alone or NPs alone, suggesting a synergistic interaction between AMF and NPs. Among interactive treatments, AMF + ZnO NPs treatment resulted in the highest overall organic acid production except citric acid which was higher produced by AMF+ Zn/Fe NPs. ZnO NPs and Zn/Fe NPs alone or in combination with AMF had mixed effects, depending on the organic acid. Overall, AMF + ZnO NPs treatment resulted in the highest production of organic acids in maize roots, as compared with control.

**Table 1:** Interactive effect of NPs and AMF on organic acids. The numbers indicate the mean value of organic acids. Numbers following means represent the standard error (± SE) of three replicates. Different letters above the bars indicate statistically significant differences among treatments (p < 0.05).

| Treatments | Oxalic (ug/g) | Citric | lactic | Malic | T-aconitic | Succinic |
| --- | --- | --- | --- | --- | --- | --- |
| Cn | 162.03±15.12d | 20.37±2.60d | 70.77±5.87b | 22.41±2.86b | 212.33±21.09a | 3.14±0.09de |
| AMF | 259.94±19.88abcd | 27.22±2.61cd | 68.59±4.63b | 39.06±2.90b | 209.08±10.39a | 2.97±0.19e |
| FeO NPs | 188.90±9.15cd | 29.06±2.56cd | 79.00±10.72b | 29.29±2.32b | 234.55±6.23a | 3.41±0.17de |
| ZnO NPs | 211.93±12.94cd | 34.07±3.56cd | 76.81±9.45b | 33.86±2.75b | 238.56±12.26a | 3.97±0.12cd |
| Zn/Fe NPs | 245.81±15.63bcd | 38.23±2.75cd | 81.07±11.38b | 38.69±2.67b | 231.68±18.45a | 4.94±0.30bc |
| AMF +FeO NPs | 295.04±26.26abc | 43.35±5.71bc | 92.37±4.49ab | 71.45±6.36a | 243.04±12.58a | 5.82±0.16ab |
| AMF+ ZnO NPs | 362.21±25.98a | 58.71±5.47ab | 121.19±5.43a | 82.62±4.78a | 260.57±16.26a | 6.31±0.34a |
| AMF+ Zn/Fe NPs | 330.15±41.60ab | 64.60±5.66a | 102.84±3.61ab | 76.72±6.77a | 251.96±19.46a | 6.13±0.12a |

### 3.7 Effects of AMF and NPs on the nutrient concentration and uptake in maize

The individual and interactive treatments of AMF and NPs increased the nutrient concentration and nutrients uptake in maize plants (Table 2 and Table S1). Among single treatments, higher N and P uptake was noted with Zn/Fe NPs, K uptake by AMF treatment, Zn uptake in ZnO NPs treatment and Fe uptake was recorded by FeO NPs, as compared with control treatment. AMF + ZnO NPs was prominent in significant *(P* ≤ *0.05)* improvement of nutrients concentration in shoot and root of maize (Table S1). Similarly, the most significant *(P* ≤ *0.05)* and higher N, P, K and Zn uptake was noted under AMF + ZnO NPs, while most significant *(P* ≤ *0.05)* and higher uptake of Fe was recorded in AMF + FeO NPs among interactive treatments, as compared with control.

**Table 2:** Interactive effects of NPs and AMF on the nutrient uptake in maize. The numbers indicate the mean value of whole plant nutrient uptake. Numbers following means represent the standard error (± SE) of three replicates. Different letters above the bars indicate statistically significant differences among treatments (p < 0.05).

| Treatments | Nutrient uptake |  |  |  |  |
| --- | --- | --- | --- | --- | --- |
|  | N (mg/plant) | P (mg/plant) | K (mg/plant) | Zn (ug/plant) | Fe (ug/plant) |
| Cn | 112.53±2.95c | 16.71±0.55d | 62.69±4.10d | 180.97±10.63d | 1838.07±106.33c |
| AMF | 197.11±32.06bc | 27.84±3.44cd | 187.25±35.32bc | 403.95±58.90bc | 4315.47±722.58bc |
| FeO NPs | 129.31±13.60c | 19.23±1.80d | 114.32±11.60cd | 272.73±26.76cd | 6331.01±544.15b |
| ZnO NPs | 164.73±9.66bc | 27.74±0.91cd | 145.50±3.51cd | 448.78±17.96bc | 2978.72±270.26c |
| Zn-Fe NPs | 204.42±10.49bc | 32.09±3.35bc | 177.87±19.24bcd | 421.82±38.84bc | 6169.92±560.12b |
| AMF +FeO NPs | 216.55±21.01bc | 32.28±2.39bc | 227.53±19.17bc | 385.30±13.27bc | 10903.21±570.41a |
| AMF+ ZnO NPs | 334.53±33.71a | 49.32±4.38c | 403.15±45.62a | 683.28±72.44a | 6729.25±911.25b |
| AMF+ Zn-Fe NPs | 262.42±37.04ab | 40.82±1.49ab | 291.85±19.91ab | 503.87±29.38ab | 11321.40±891.82a |

### 3.8 Relationship and correlations revealed association among various traits

The partial least squares path model (PLS-PM) results showed that relationships were observed between plant growth parameters, phosphatase activity, rhizosheath pH, mycorrhizal efficiency, organic acids, nutrient uptake (Fig. 11). Among them, mycorrhizal efficiency showed positive correlations with organic acids (path coefficient: 0.842), nutrients uptake (path coefficient: 0.367), while a negative correlation was observed with rhizosheath pH (path coefficient: −0.227). Rhizosheath pH showed positive associations with organic acids (path coefficient: 0.037), nutrient uptake (path coefficient: 0.263), and root growth (path coefficient: 11.840), indicating its role as a mediating factor in nutrient availability and plant growth. However, organic acids displayed positive correlations with root growth (path coefficient: 52.582) and nutrient uptake (path coefficient: 0.710), but they inversely affected phosphatase activity (path coefficient: − 0.544). Moreover, nutrient uptake exhibited positive correlations with phosphatase activity (path coefficient: 1.463) and shoot growth (path coefficient: 0.994), while having a strong negative impact on root growth (path coefficient: −51.161). Root growth also negatively influenced shoot growth (path coefficient: −0.030), indicating complex trade-offs in biomass allocation under varying nutrient availability.

**Figure 11:**
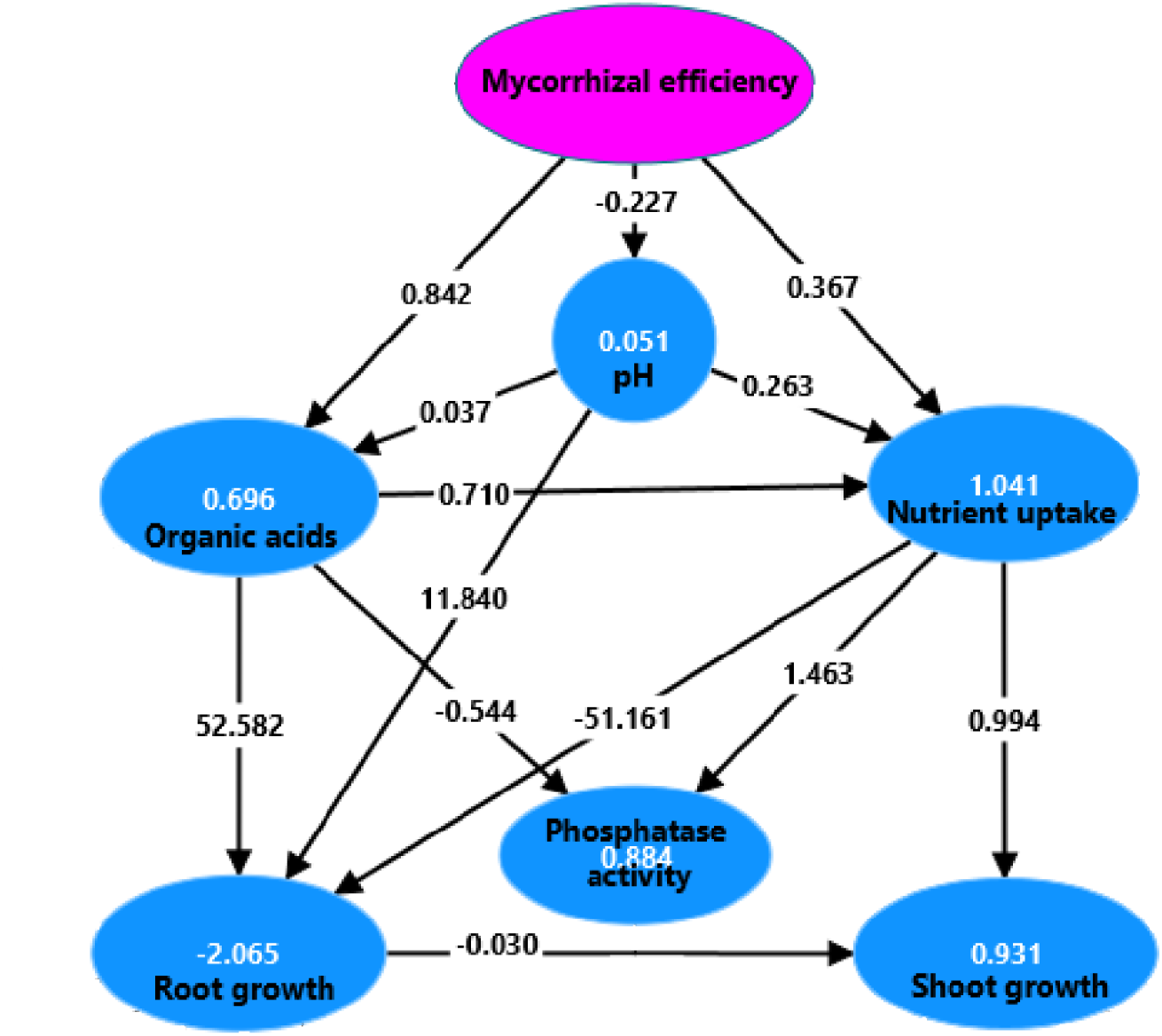
Partial least squares path model (PLS-PM) illustrating the relationships between nutrient uptake, root system architecture, shoot growth traits, organic acids, rhizosphere pH, phosphatase activity, and mycorrhizal efficiency in maize. The values within the latent variable circles represent coefficients of determination (*R*^2^), indicating the proportion of variance explained by the model for each construct. Black arrows indicate directional relationships between latent variables, and the values along the arrows represent the path coefficients, denoting the strength and positive or negative influence of these relationships. High path coefficients indicate stronger influences between constructs.

The Pearson correlation analysis further validated the associations among the traits of maize treated with NPs and AMF (Fig. 12). Strong positive correlations were found between shoot and root growth parameters, rhizosphere pH, phosphatase activity, organic acids, and nutrient uptake, suggesting a complementary interaction of NPs and AMF in promoting maize growth. Improvements in one growth aspect, such as root or shoot biomass, were generally accompanied by enhancements in nutrient uptake and other physiological traits. Interestingly, the correlation matrix did not reveal any strong negative relationships, highlighting a general synergy among the measured traits. pH, however, exhibited only weak correlations with organic acids, implying that despite the role of these acids in nutrient solubilization, their presence did not significantly alter rhizosheath pH under the given experimental conditions. The findings from both analyses underscore the potential of nutrient management in optimizing plant growth and the importance of organic acids in modulating complex biochemical interactions within the rhizosphere.

**Figure 12:**
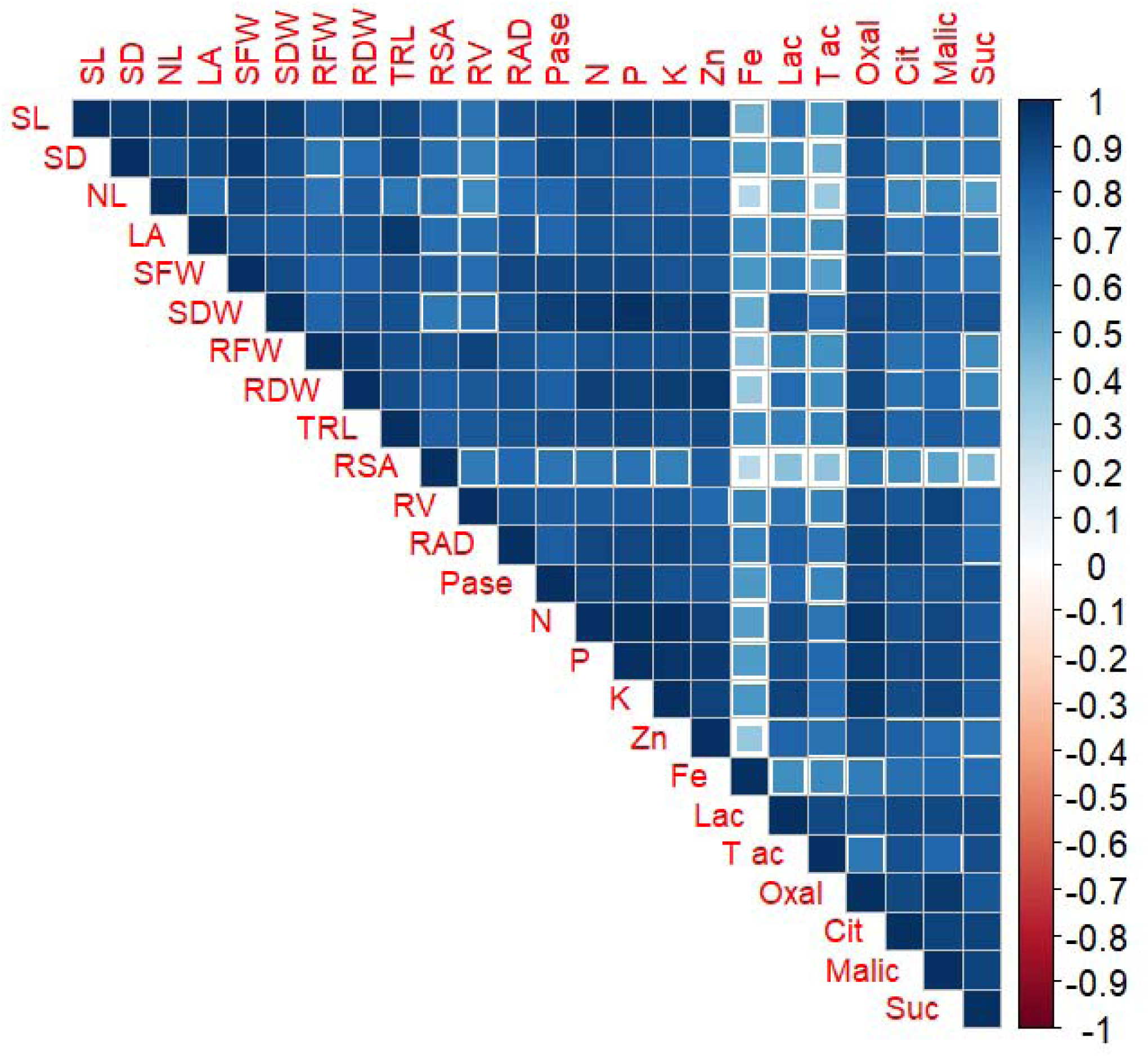
Pearson’s correlation among traits of maize under the treatments of NPs and AMF. The color gradient represents correlation coefficients, with blue indicating positive correlations (closer to 1), white indicating no correlation (0), and red indicating negative correlations (closer to −1). Trait notations: shoot length (SL), shoot diameter (SD), number of leaves (NL), leaf area (LA), shoot fresh weight (SFW), shoot dry weight (SDW), root fresh weight (RFW), root dry weight (RDW), total root length (TRL), root surface area (RSA), root volume (RV), and root architecture diameter (RAD), phosphatase activity (Pase), rhizosheath pH (pH), nitrogen (N), phosphorus (P), potassium (K), zinc (Zn), and iron (Fe), lactic acid (Lac), T-aconitic acid (T-ac), oxalic acid (Oxal), citric acid (Cit), malic acid (Malic), and succinic acid (Suc).

The strong correlations between growth parameters and nutrient uptake emphasize the need for targeted strategies to enhance nutrient assimilation and plant development. Furthermore, the observed diverse relationships involving organic acids highlight the complexity of the underlying metabolic networks, warranting further experimental investigations to elucidate the mechanistic pathways involved.

## 4. Discussion

Applying NPs and AMF together offers a strong way to increase plant production and development. AMF enhance plant nutrient absorption, water uptake, and resilience against environmental stressors. Concurrently, NPs can amplify these advantages by optimizing nutrient delivery systems, modifying soil chemistry, and elevating plant stress tolerance. The synergistic effects of AM fungus and green NPs mediated by *Moringa oleifera* on maize growth, functional characteristics of the roots, and nutrient uptake were investigated in this work.

The results revealed that NPs and AMF, individually and in combination, significantly (*P* ≤ *0.05*) enhanced maize shoot and root growth parameters (Fig. 8). The improved above and below ground traits of maize under these treatments confirms previous findings on the role of NPs and AM symbiosis in improving plant growth, nutrient acquisition, and stress resilience (El-Gazzar et al., 2020; Naseer et al., 2022). However, It was discovered that utilizing zinc oxide and selenium NPs in conjunction with AMF offered tactical benefits for enhancing the growth and yield of chili plants under cold stress circumstances (Sayed et al., 2024). This enhancement is likely due to the distinct characteristics of NPs and AMF; notably, mycorrhizae form extensive networks in the soil that effectively extend the root system’s reach. This allows for more efficient water and nutrient absorption, a critical factor in environments with limited resource availability (Huey et al., 2020; Alaux et al., 2021). However, NPs improved soil penetration and nutrient solubilization, which can directly affect nutrient availability to the plant roots (ul Ain et al., 2023), and these NPs function as micro-reservoirs of nutrients that slowly release ions into the soil, which are then readily available for uptake by the mycorrhizal network.

Our results demonstrate that ZnO, Zn/Fe NPs, and AMF, either independently or in combination, significantly enhance phosphatase activity (Fig. 9). These findings indicate that AMF, ZnO, and Zn/Fe NPs improve phosphatase activity through nutrient enhancement, protective antioxidant properties, and direct enzyme activation. Interestingly, while the application of FeO NPs alone reduced phosphatase activity, this effect was reversed when FeO NPs were used in conjunction with AMF (AMF + FeO NPs). FeO NPs alone appear to decrease phosphatase activity, likely due to induced oxidative stress and enzyme inhibition (Cao et al., 2017; Cameron et al., 2022). However, the detrimental effects of FeO NPs were ameliorated when these NPs were applied in combination with AMF, which likely provides a synergistic protective effect, thereby improving enzyme activity (Feng et al., 2013; Cao et al., 2016).

Our results indicated that ZnO NPs were the most effective in increasing root colonization whereas FeO, Zn/Fe NPs reduced it. Interestingly, ZnO and Zn/Fe NPs were found to enhance mycorrhizal dependency, whereas FeO NPs had a reducing effect (Fig. 10). Supporting these observations, previous studies have reported that 400 mg/kg dose of ZnO NPs increase root colonization (Wang et al., 2016). In contrast, nanoscale zerovalent iron (Yang et al., 2024), and zinc ferrite NPs have been reported to decrease root colonization (Metwally and Abdelhameed, 2024). The effects of NPs on mycorrhizal colonization and dependency varies, influenced by the specific NPs type, with some NPs enhance colonization while others inhibit it (Cao et al., 2016; Cao et al., 2017; Tian et al., 2019; Naseer et al., 2022; Sayed et al., 2024).

Our findings indicated that the interaction between AMF and green-synthesized NPs substantially impacts the production of organic acids in maize roots. While individual treatments with AMF and NPs variably influenced specific acids, their combined application generally resulted in a synergistic increase in organic acid levels, with the AMF + ZnO NPs treatment yielding the enhancement across most organic acids measured (Table 1). However, individual and interactive treatments of AMF and NPs markedly enhanced nutrient concentration and uptake in maize, with interactive treatments showing the greatest impact (Table 2). The NPs enhance nutrient absorption by increasing nutrient availability and solubility in the rhizosphere through their high surface area and reactivity, which facilitate improved nutrient uptake efficiency (Singh et al., 2024). For instance, ZnO and FeO NPs promote the release of zinc and iron ions, respectively, which are essential for plant metabolic activities (Chandrika et al., 2022; ul Ain et al., 2023). On the other hand, AMF improve nutrient absorption by expanding the root surface area through their extensive hyphal networks, increasing nutrient acquisition from a larger soil volume, and enhancing root nutrient transporter activities (Hussain et al., 2021; Ain et al., 2024). In combined treatments, NPs and AMF synergistically enhance nutrient uptake (Ostadi et al., 2022; Sayed et al., 2024) by concurrently improving nutrient availability by NPs (ul Ain et al., 2023) and increasing root absorption capacity by AMF (Huey et al., 2020), resulting in superior nutrient acquisition compared to individual applications (Ostadi et al., 2022).

## 5. Conclusion

The combined application of Moringa oleifera-mediated green nanoparticles (NPs) and arbuscular mycorrhizal fungi (AMF) has emerged as a promising strategy to enhance maize growth, root functional traits, root exudation, mycorrhizal colonization, and nutrient uptake. Fig. 13 presents a schematic diagram summarizing the synergistic effects of Moringa oleifera mediated green NPs and AMF on maize plants. Among the individual treatments, Zn doped FeO NPs demonstrated the greatest improvement in maize growth compared to other nanoparticle treatments. However, in combination with AMF, ZnO NPs exhibited superior performance over both FeO and Zn doped FeO NPs in promoting overall plant growth and development. These findings underscore the potential of environmentally friendly, plant-mediated NPs as a sustainable agricultural tool to enhance crop resilience and productivity, particularly in stress-prone environments. Moving forward, future research should emphasize long-term field trials and investigate the ecological impacts of green NPs and AMF on soil health and plant systems to establish their safety and efficacy in sustainable agricultural practices.

**Figure 13:**
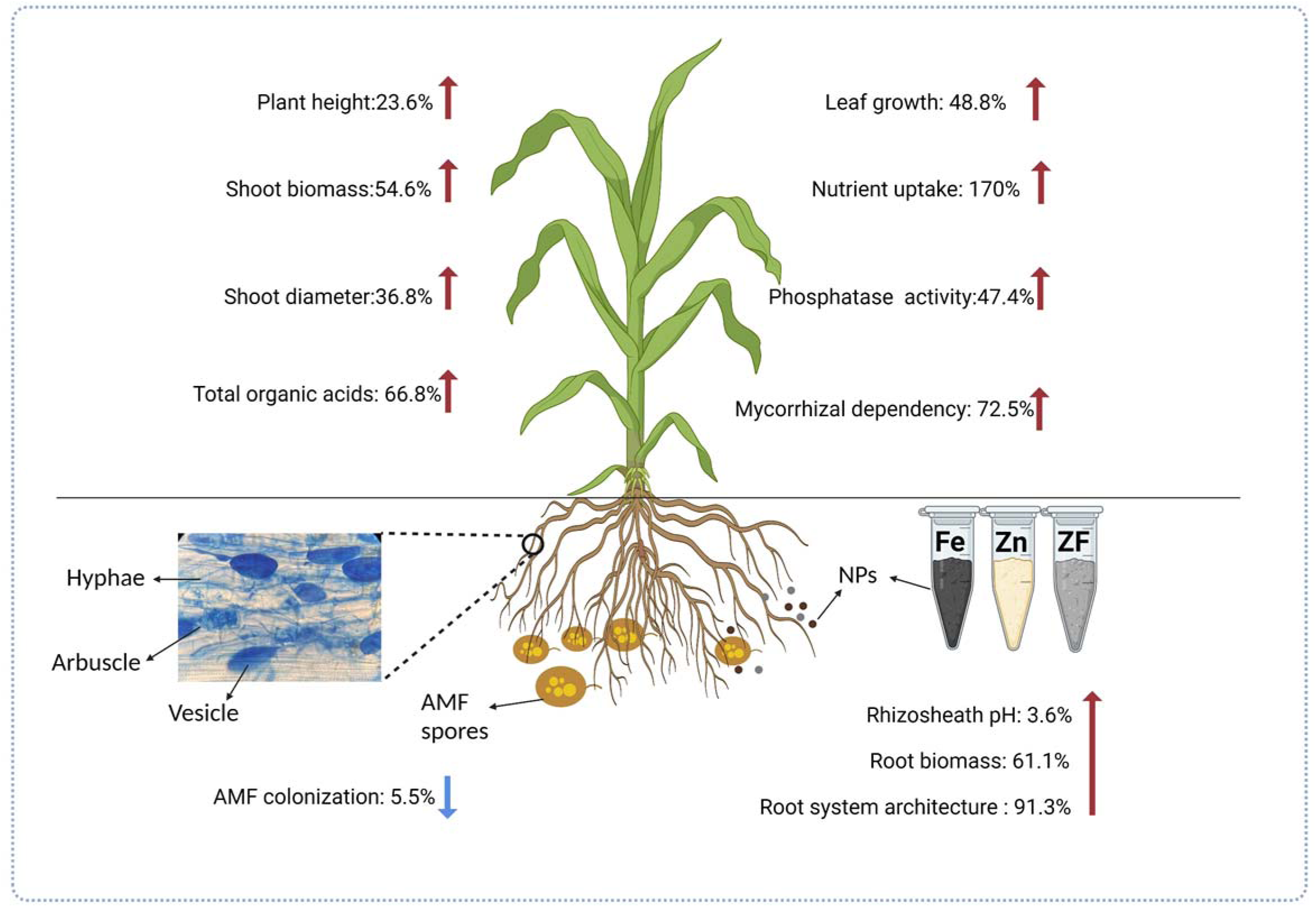
Systematic illustration of interactive effects of *moringa oleifera* mediated green NPs and arbuscular mycorrhizal fungus (AMF) on growth, root functional traits, and nutrient uptake of maize (Zea mays L.). Red arrows indicate increases and blue arrows show a decrease in growth traits.

