## supplementary material for "Interactive effect of *Moringa oleifera* mediated green nanoparticles and arbuscular mycorrhizal fungi on growth, root system architecture, and nutrient uptake in maize (*Zea mays* L.)"

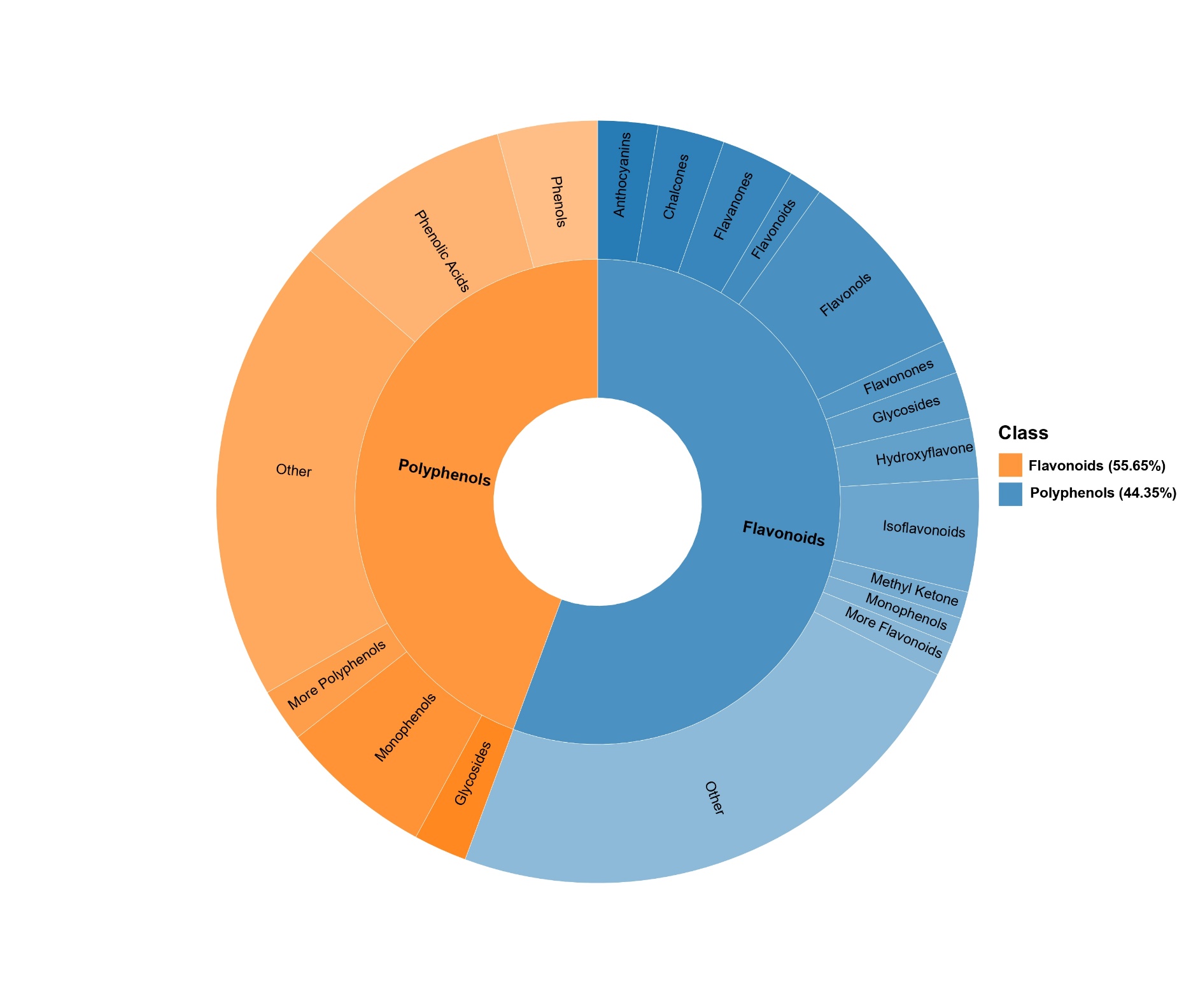

**Fig S1:** Sunburst diagram representing the composition of metabolite classes. Each color denotes a distinct metabolite category, with the area of each section indicating its relative proportion. The inner circle represents the primary classification of metabolites, while the outer circle provides a more detailed secondary classification. Flavonoids (blue) account for 55.65% of the total composition, whereas polyphenols (orange) make up 44.35%.

**Table S1:** Interactive effects of NPs and AMF on the nutrient concentration in maize shoot and root. The numbers indicate the mean value of organic acids. Numbers following means represent the standard error (± SE) of three replicates. Different letters above the bars indicate statistically significant differences among treatments (p < 0.05).

| **Treatments** | **Shoot** | | | | | **Root** | | | | |
| --- | --- | --- | --- | --- | --- | --- | --- | --- | --- | --- |
|  | **N (mg/g)** | **P (mg/g)** | **K (mg/g)** | **Zn (ug/g)** | **Fe (ug/g)** | **N (mg/g)** | **P (mg/g)** | **K (mg/g)** | **Zn (ug/g)** | **Fe (ug/g)** |
| **Cn** | 19.10±1.33a | 2.88±0.24b | 10.20±1.20e | 21.73±1.45c | 150.57±16.04e | 22.50±2.42a | 3.21±0.16d | 14.16±1.05c | 64.70±3.79a | 890.70±46.24c |
| **AMF** | 24.90±1.59a | 3.38±0.10ab | 22.60±2.46bcd | 43.42±2.25ab | 331.20±23.36de | 26.40±2.08a | 4.19±0.22bcd | 27.40±2.76bc | 75.70±5.30aa | 1162.10±116.19c |
| **FeO NPs** | 19.53±1.70a | 3.01±0.23ab | 14.70±1.22de | 35.41±2.59bc | 555.60±26.49ab | 24.20±2.47a | 3.40±0.21cd | 29.70±2.89b | 69.60±5.51a | 2419.70±225.92b |
| **ZnO NPs** | 20.50±2.03a | 3.54±0.21ab | 17.43±1.86cde | 53.23±5.09a | 147.93±16.36e | 25.10±2.34a | 3.88±0.29bcd | 24.10±2.71bc | 76.17±2.19a | 1104.67±114.72c |
| **Zn/Fe NPs** | 22.60±1.76a | 3.66±0.27ab | 19.12±1.51cde | 41.75±1.73ab | 535.10±22.57bc | 27.80±3.56a | 3.96±0.28bcd | 26.60±2.82bc | 76.20±5.35a | 1393.10±51.18c |
| **AMF +FeO NPs** | 25.80±2.50a | 3.77±0.27ab | 26.20±2.10abc | 39.81±2.65ab | 725.80±70.97a | 29.57±3.58a | 4.62±0.31abc | 33.80±1.97ab | 73.10±7.65a | 3348.70±242.22a |
| **AMF+ ZnO NPs** | 27.60±1.84a | 4.03±0.12a | 32.10±2.08a | 53.64±3.22a | 370.60±43.64cd | 34.90±2.04a | 5.21±0.32ab | 45.10±4.59a | 80.40±7.65a | 1309.60±152.93c |
| **AMF+ Zn/Fe NPs** | 26.20±3.15a | 3.96±0.16a | 29.37±2.20ab | 46.80±3.44ab | 561.70±45.88ab | 30.77±1.67a | 5.59±0.37a | 36.70±3.31ab | 77.10±3.22a | 3331.10±182.70a |

**Materials and Methods**

**Detailed methodology of metabolic analysis**

The plant polyphenols metabolomics analysis was conducted using a UPLC-ESI-MS/MS system (UPLC: Waters Acquity I-Class PLUS; MS: Applied Biosystems QTRAP 6500+). Sample preparation involved freeze-drying the samples in a vacuum, followed by weighing 50 mg of the sample and extracting it with 1000 μL of a methanol:acetonitrile:water solution (1:2:1, v/v/v). The samples were vortexed for 30 seconds, ground with steel balls at 45 Hz for 10 minutes, and ultrasonicated in an ice-water bath for 10 minutes. After incubation at -20°C for one hour, the samples were centrifuged at 4°C at 12,000 rpm for 15 minutes. A 300 μL aliquot of the supernatant was carefully collected and filtered through a 0.22 μm organic filter membrane into a 2 mL injection vial. Quality control (QC) samples were prepared by pooling 10 μL from each sample. Chromatographic separation was performed on a Waters HSS-T3 column (1.8 µm, 2.1 mm × 100 mm) with a gradient mobile phase consisting of solvent A (0.1% formic acid and 5 mM ammonium acetate in water) and solvent B (0.1% formic acid in acetonitrile). The flow rate was set at 0.35 mL/min, and the column oven was maintained at 50°C. The injection volume was 4 μL. The gradient program spanned 13 minutes, beginning with 98% A and 2% B, with progressive adjustments to achieve optimal separation. Mass spectrometry was performed using an ESI-triple quadrupole-linear ion trap (QTRAP)-MS with parameters optimized for sensitive detection. Identified compounds were annotated for classification and pathway mapping using KEGG, HMDB, and LipidMaps databases.

For targeted metabolome analysis, 23.3 mg of each sample was weighed into a 2 mL centrifuge tube, homogenized with steel balls, and extracted with 1 mL of a methanol:water solution (7:3, v/v) containing internal standards. After vortexing for 30 seconds, the samples were homogenized at 45 Hz for 15 minutes and ultrasonicated in an ice-water bath for 30 minutes. Centrifugation at 4°C and 12,000 rpm for 15 minutes yielded a supernatant, of which 200 μL was filtered through a 0.22 μm membrane for analysis. Chromatographic separation was achieved using a Waters ACQUITY I-Class system equipped with an HSS T3 column (100 × 2.1 mm, 1.8 μm). The mobile phase consisted of solvent A (0.1% formic acid in water) and solvent B (0.1% formic acid in acetonitrile), with the column and auto-sampler temperatures set at 35°C and 10°C, respectively. A SCIEX QTRAP 6500+ triple quadrupole mass spectrometer, fitted with an IonDrive Turbo V electrospray ionization (ESI) interface, was used for detection. The instrument parameters, including curtain gas (35 psi), ion spray voltage (+5500 V/-4500 V), source temperature (550°C), and gas pressures, were optimized for sensitivity and precision. Multiple reaction monitoring (MRM) transitions were optimized for each analyte, with the most sensitive Q1/Q3 transitions used for quantification and validation. Data acquisition and processing were performed using SCIEX Analyst Workstation Software (v1.7.2) and Sciex OS 2.0.1. Calibration curves were constructed using analyte concentrations ranging from 10 nmol/L to lower limits, with linear regression and 1/x weighting ensuring high accuracy within 80–120%. The limits of detection (LOD) and quantitation (LOQ) were determined based on signal-to-noise ratios of 3 and 10, respectively. Precision was assessed by determining the relative standard deviation (RSD) of replicate QC sample injections, while accuracy was evaluated by calculating the percent recovery of spiked QC samples.
